# SEISMICgraph: a web-based tool for RNA structure data visualization

**DOI:** 10.1101/2024.09.26.615187

**Authors:** Federico Fuchs Wightman, Grant Yang, Yves J. Martin des Taillades, Casper L’Esperance-Kerckhoff, Scott Grote, Matthew F. Allan, Daniel Herschlag, Silvi Rouskin, Lauren D. Hagler

## Abstract

In recent years, RNA has been increasingly recognized for its essential roles in biology, functioning not only as a carrier of genetic information but also as a dynamic regulator of gene expression through its interactions with other RNAs, proteins, and itself. Advances in chemical probing techniques have significantly enhanced our ability to identify RNA secondary structures and understand their regulatory roles. These developments, alongside improvements in experimental design and data processing, have greatly increased the resolution and throughput of structural analyses. Here, we introduce SEISMICgraph, a web-based tool designed to support RNA structure research by offering data visualization and analysis capabilities for a variety of chemical probing modalities. SEISMICgraph enables simultaneous comparison of data across different sequences and experimental conditions through a user-friendly interface that requires no programming expertise. We demonstrate its utility by investigating known and putative riboswitches and exploring how RNA modifications influence their structure and binding. SEISMICgraph’s ability to rapidly visualize adenine-dependent structural changes and assess the impact of pseudouridylation on these transitions provides novel insights and establishes a roadmap for numerous future applications.

**GRAPHICAL ABSTRACT:** 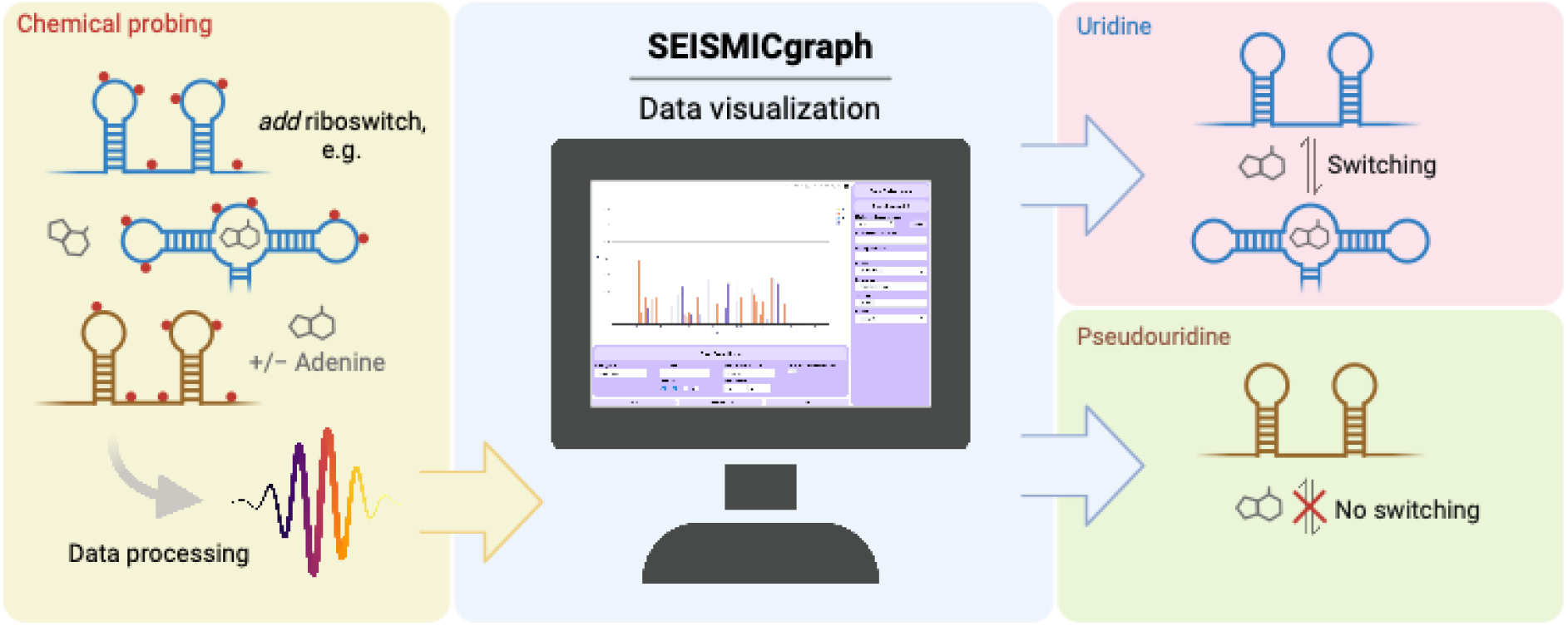

## INTRODUCTION

Since its initial discovery as an intermediary of genetic information, RNA has been recognized for its diverse regulatory roles, including the control of transcription, translation, splice site selection, and RNA degradation (Allan et al., 2023; Wan et al., 2011). These functions are achieved through RNA’s ability to bind proteins and fold into complex structures. The discovery of novel RNA interactions and their biological function has driven and been supported by the development of numerous techniques, each with specific advantages and limitations (Jones, 2016; Ma et al., 2022; Spitale and Incarnato, 2023; Turner and Mathews, 2016; Zhang et al., 2022).

Chemical probing methods provide detailed insights into RNA base-pairing, folding, and interactions with biomolecules such as proteins and ligands at single-nucleotide resolution. These techniques use small molecules such as dimethyl sulfate (DMS) and SHAPE reagents (e.g., 1-methyl-7-nitroisatoic anhydride (1M7)) to specifically and covalently modify RNA at chemically accessible sites (Ehresmann et al., 1987; Merino et al., 2005; Peattie and Gilbert, 1980; Soukup and Breaker, 1999). The differential chemical reactivity between nucleotides is used to distinguish between folded and unfolded regions of a given RNA structure (Douds et al., 2024; Siegfried et al., 2014; Zubradt et al., 2017). DMS methylates the Watson-Crick-Franklin face of solvent-exposed and unpaired A and C bases, while SHAPE reagents selectively acylate the 2’-hydroxyl group of bases in non-based paired regions that can attain the reactive configuration (Lucks et al., 2011; Rouskin et al., 2014; Watts et al., 2009). Both DMS and SHAPE techniques can be combined with mutational profiling (MaP), where chemically modified residues are converted into mutations during reverse transcription to enable the generation of high-resolution structural maps across RNA molecules (Homan et al., 2014; Siegfried et al., 2014; Zubradt et al., 2017). These methods are highly adaptable and can be employed to study how environmental conditions impact RNA interactions, allowing comparisons between *in vitro* and cellular contexts and providing new insights into the relationship between RNA sequence, structure, and function.

As chemical probing methods have become more widely adopted, several computational tools have been developed to analyze the resulting sequencing data. These include DREEM (Tomezsko et al., 2020), RNA Framework (Incarnato et al., 2018), and ShapeMapper 2 (Busan and Weeks, 2018), and the more recent SEISMIC-RNA (Allan et al., 2024), which builds upon DREEM. These tools facilitate the alignment of sequencing reads to reference sequences, the identification and quantification of mutations, and the generation of mutation profiles for each reference. The data generated through these analyses can be further used to test hypothesis-driven models and draw biophysical or biological conclusions, including the generation of secondary structure models and the assessment of the effects of mutations or environmental changes. Despite their utility, these tools typically require users to have at least basic programming skills to perform quality control assessments, visualize data, and compare samples, posing challenges for those less familiar with bioinformatics.

To overcome these limitations and allow the broader community to effectively use these approaches, we introduce SEISMICgraph, a web-based platform. SEISMICgraph is designed to streamline and standardize the visualization and analysis of chemical probing data. It was validated using datasets from DMS-MaPseq experiments involving well-characterized RNA structures. The application provides an accessible framework for generating plots and analyzing raw chemical probing data from RNA libraries. SEISMICgraph enables researchers to produce standard visualizations of chemical probing data, making the interpretation of results more accessible and interpretable, while also facilitating broad use in the scientific community. SEISMICgraph is available at https://seismicrna.org.

## MATERIALS AND METHODS

### Library Design

The library used for the initial analysis contained 48 RNA structures and uses a common primer, flank, and barcoding strategy (Figure 2A). Library amplification, in vitro transcription, DMS-MaPseq, and data processing was performed as outlined below with minor changes. Specifically, DMS reactions varied in magnesium (MgCl_2_) concentration (0, 1, and 10 mM), while keeping the RNA input (10 μg), buffer (150 mM potassium cacodylate, pH 7.4), DMS concentration (5% v/v), reaction time (5 min), and temperature (37 °C) constant. One reference (*add* aptamer domain) and section (variable region, bases 40-112) were chosen from the library to help visualize the plot types in Figure 2.

**Figure 1.**
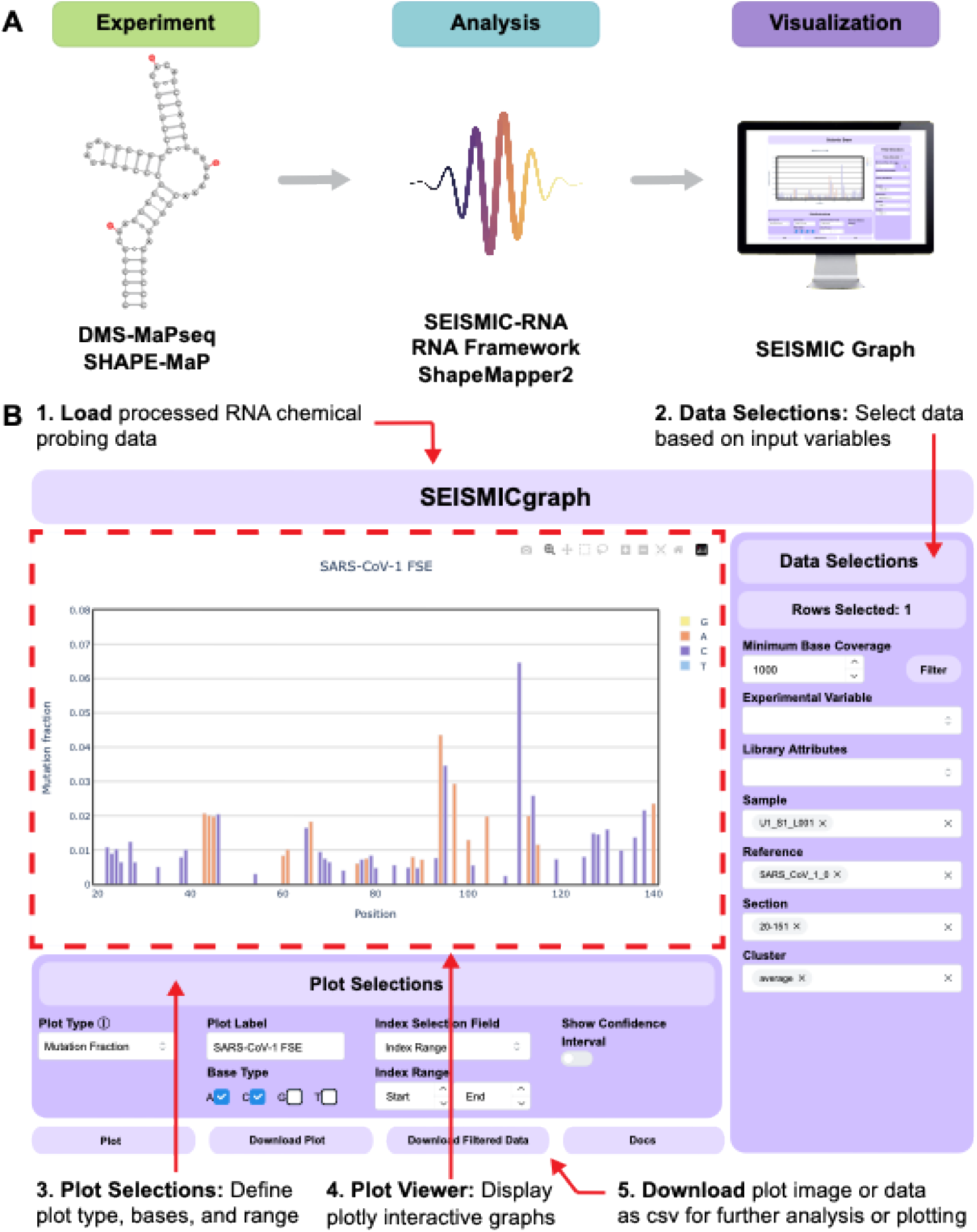
SEISMICgraph. (A) To study RNA secondary structures, DMS-MaPseq or SHAPE-MaP is performed by chemically modifying unstructured regions of RNA depicted here with red circles. Modifications are converted to mutations during RT-PCR and read out by next generation sequencing. The sequencing data from these experiments is then analyzed with SEISMIC-RNA, RNA Framework, or ShapeMapper 2 before being visualized with SEISMICgraph (see also Supplementary Figure 1). (B) Overview of the SEISMICgraph interface.

**Figure 2.**
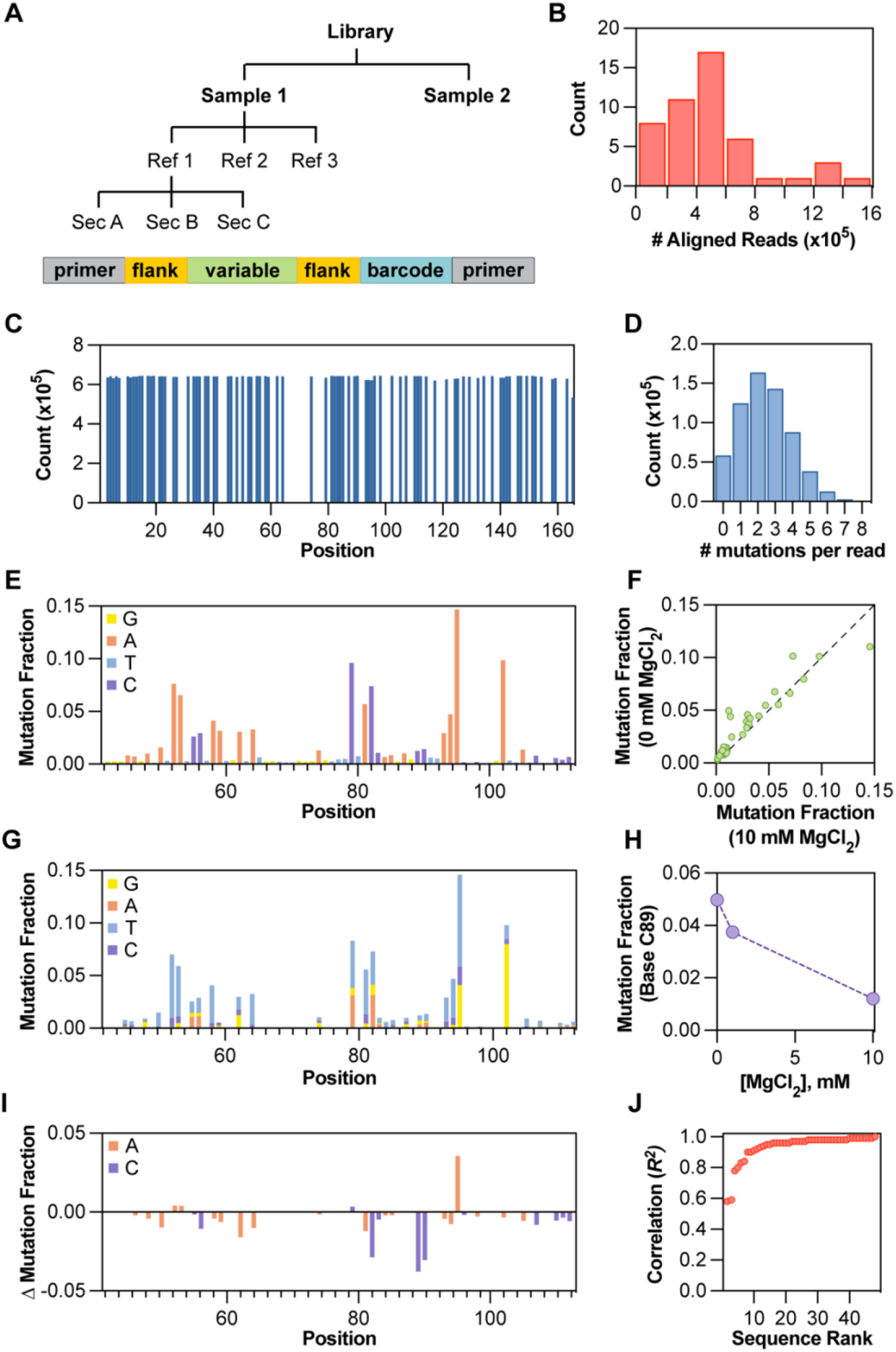
Examples of plot types available in SEISMICgraph. (A) Hierarchical data structure used in SEISMICgraph. For this library, each sample was separated into its reference sequences, which in turn were divided into sections. The references were structured to have common primers for library-specific amplification, a reference-specific barcode, variable-length flanking sequences to ensure all library members are of the same size, and a variable region that contains the RNA sequence of interest (see Methods and Supplementary Information). (B) **# Aligned reads / Reference as Freq. Dist.**for one sample of a designed library of known RNA structures. (C-I) Example plots for one reference in the library where the variable region of interest is the *add* riboswitch (*Vibrio vulnificus*) aptamer domain. (C) **Base coverage** for entire amplicon (showing only As and Cs, section = full). (D) **Mutation per Read per Reference** shows the frequency distribution of mutations per read in bins. (E) **Mutation Fraction** for the variable section. Bases are colored by their identity in the reference sequence where A is orange, C is purple, T is light blue, and G is yellow. (F) **Compare Mutation Profiles** for two samples with no MgCl_2_ or 10 mM MgCl_2_. Each green circle represents one reactive A or C base (*N* = 33; Pearson *R* = 0.93, RMSE = 0.011). The black dashed line represents the line of identity predicted for the null model of no difference between the samples (x = y). (G) **Mutation Fraction Identity** for the variable section. Each stacked bar represents the fraction mutated to that base, where orange represents bases mutated to A, purple represents bases mutated to C, light blue represents bases mutated to T, and yellow represents bases mutated to G. (H) **Experimental Variable Across Samples**. The mutation fraction from 3 samples (0, 1, and 10 mM MgCl_2_) are shown for one base (C at position 89). (I) **Mutation Fraction Delta** comparing the variable section of two samples (0 and 10 mM MgCl_2_) for only A and C bases. The change in mutation fraction here is calculated as sample 1 (10 mM MgCl_2_) – sample 2 (0 mM MgCl_2_). Bases are colored by identity in the reference where A is orange and C is purple. (J) **Correlation by Refs. Between Samples**. The Pearson correlations (*R*^2^) for all the members of the library were calculated by comparing the 0 vs. 10 mM MgCl_2_ samples, where each red circle is an individual reference sequence.

A second library had the same design strategy (Figure 3A). We retrieved from the RiboswitchDB database (Mukherjee et al., 2019) 29 predicted Mg^2+^-dependent riboswitches and 30 proposed to respond to purines. We included the *add* riboswitch from *Vibrio vulnificus* as a positive control. The total library encompasses 81 sequences, including above mentioned putative riboswitches and other well-known structures such as the HIV FSE (see Supplementary Table 1). To enable pooled chemical probing with DMS-MaPseq, we designed the library to include the following: (1) Common forward and reverse primer sequences (forward primer: 5’-TTAAACCGGCCAACATACC -3’; reverse primer: 5’-TCGAAAGGAACGAGTAGCG -3’). (2) Variable length flanking sequences to make all the constructs of equal length. Flanking sequences were generated by randomly shuffling the order of Cs and Us. Sequences were filtered for consecutive stretches of Cs or Us longer than 3 nucleotides and were regenerated for these cases. (3) Unique barcodes were added to each sequence for later demultiplexing (Hamming distance > 3), which was required because of similarities between several library sequences and to identify parent sequences after DMS modification (Figure 2A and 3A). In combination, these features increase sequencing read depth and quality, by decreasing RT and/or PCR biases (thereby giving more uniform reads of all library members) and by unambiguously assigning very similar sequences in the library pool. The complete sequences of the 81-member library are given in Supplementary Table 1. The DNA library was obtained as an oligo pool from Integrated DNA Technologies (IDT).

**Figure 3.**
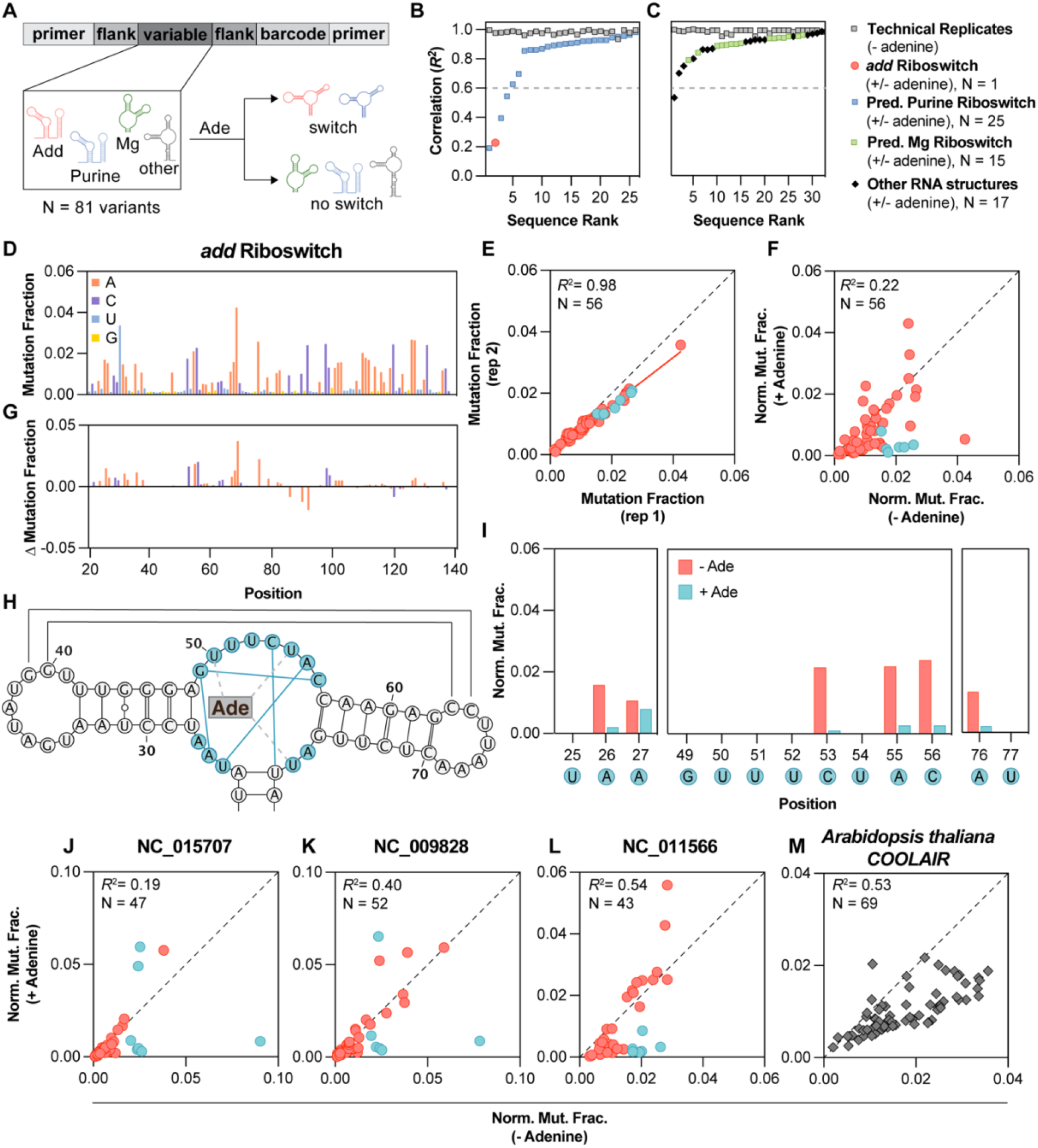
Experimental identification of adenine-responsive RNAs. (A) Library design for the riboswitch library. In this experiment the structure is probed with DMS in the presence or absence of adenine (Ade) to assess which structures undergo a conformation change. (B-C) Ranked correlation of references compared to each other in technical replicates (gray squares) or in samples with and without 5 mM adenine. (D) Mutation fraction at each position of the riboswitch section. (E) Comparison of mutation fraction at each position of two technical replicates where each red circle is an A or C base (N = 56). Turquoise circles depict A and C bases in the aptamer binding domain as in H. The Pearson correlation (*R*^*2*^) was calculated in SEISMICgraph. The black dashed line represents the line of identity, and the red solid line represents the best-fit line from linear regression (slope = 0.77). (F) Comparison of the mutation fraction of samples in the presence and absence of 5 mM adenine. (G) Mutation fraction delta comparing samples in the presence and absence of 5 mM adenine, where the difference is calculated as sample 1 (with adenine) – sample 2 (without adenine). (H) Predicted secondary structure of the aptamer domain of the *add* riboswitch (Tian et al., 2018), where bases in the adenine binding site are depicted as turquoise circles. Hydrogen bond interactions enforced by binding are between bases in the binding site are shown with turquoise solid lines (Serganov et al., 2004). Hydrogen bonds between the binding site and adenine are shown as grey dashed lines (Serganov et al., 2004). (I) Comparison of bases in the binding site in the absence (red bars) and presence (turquoise bars) of 5 mM adenine. (J-L) Putative purine riboswitches that undergo conformational change in response to adenine. Bases are colored as in (F). (M) A sequence from *A. thaliana* that has a significant change in correlation upon addition of adenine. For (F, G, and I-M) samples were normalized as described in the Supplementary Information. See also Supplementary Figures 2, 4, and 5.

### DMS-MaPseq

The DNA library was first amplified via PCR, adding a T7 promoter sequence at the forward primer (5′-TAATACGACTCACTATAG -3′), using the Q5^®^ High-Fidelity 2X Master Mix (NEB, Cat #M0492L) and the following program: 98 °C for 2 minutes; 6 cycles of 98 °C for 10 seconds, 63 °C for 30 seconds and 72 °C for 30 seconds; and a 2 minute final extension at 72 °C and final hold at 4 °C (and for long term storage, stored at -20 °C). The annealing temperature was calculated using the NEB’s Tm calculator (https://tmcalculator.neb.com/#!/main). The reaction was carried in a final volume of 50 µl, adding 2.5 µl of each primer (10 µM) and 1 µl of the oligo pool (0.1 pmol/µl). The cycle number was optimized by screening increasing numbers of cycles until non-specific amplification was observed. To obtain the necessary yield of PCR product for the downstream applications, several reactions were run in parallel, pooled and purified using a Zymo DNA Clean & Concentrator kit (Cat. D4004).

*In vitro* transcription was performed using the T7 MEGAshortscript kit (Thermo Fisher, Cat. AM1354), starting with 200 ng of the amplified library. 20 µl reactions were incubated for 2-3 hours, followed by DNase treatment for 15 minutes. For pseudouridine experiments, UTP was replaced with 75 mM ΨTP or m1ΨTP. A yield of 30-50 µg was obtained per reaction. The RNAs were purified using the Zymo RNA Large Sample Clean-Up Kit (Cat. R1017). For chemical probing, 5-10 µg of the purified RNAs were diluted in 10 µl of nuclease-free water, denatured for 1 minute at 95 °C, and placed on ice for 2 minutes. The RNA samples were refolded in 300 mM sodium cacodylate, pH 7.2, 6 mM MgCl_2_, and optionally 5 mM adenine (or an equivalent volume of DMSO as a control). Refolding buffer (85 µl) was added to the RNAs, which were then incubated at 37 °C for 20 minutes in a thermomixer with a heated lid. Afterward, 5 µl of DMS (dimethyl sulfate) was added to the RNA solution and the reaction was allowed to proceed at 37 °C for 5 minutes with 800 rpm shaking in a thermomixer. The reaction was quenched by adding 60 µl of β-mercaptoethanol. The DMS-treated RNAs were purified using the Zymo RNA Clean & Concentrator-5 Kit (Cat. R1013). As recoveries can be as low as 10-20%, increasing the initial quantity of RNA per reaction is sometimes helpful.

The purified RNAs were reverse transcribed using the Induro Reverse Transcriptase (NEB #M0681), as described previously with minor modifications (Romero-Agosto et al., 2023). The reaction was performed as follows: 500-1000 ng of RNA, 1 µl of the reverse primer at 10 µM (sequence: 5’ TCGAAAGGAACGAGTAGCG -3’), 1 µl of 10 mM dNTPs, 0.2 µl of RNase Inhibitor, Murine (40 U/µl), 4 µl of the 5X Induro RT buffer, 1 µl of Induro Reverse Transcriptase (200 U/µl), and nuclease-free water up to 20 µl; the program was set for 30 minutes at 60 °C, followed by the addition of 1 µl of 4 M NaOH for 3 minutes at 95 °C. The cDNAs were purified using the Zymo Oligo Clean & Concentrator kit (Cat# D4060) and gave a yield of approximately 250-500 ng. We then amplified the constructs via PCR using the Q5^®^ High-Fidelity 2X Master Mix (NEB, Cat #M0492L) in a final volume of 50 µl and the following program: 98 °C for 2 minutes, 15 cycles of 98 °C for 10 seconds, 63 °C for 30 seconds and 72 °C for 30 seconds, with a 2-minute final extension at 72 °C and a final hold at 4 °C.

Samples were indexed using NEBNext Ultra II DNA Library Prep Kit for Illumina (#E7645S/L), following the manufacturer’s instructions, and sequenced on an Illumina NextSeq 1000 using a P1 (2×150 read) cartridge (Cat# 20050264).

### SEISMIC-RNA Data Processing

The first round of demultiplexing (separating the reads corresponding to each sample by their library index) was automatically performed by the Illumina platform BaseSpace (BCL-Convert 2.4.0) upon completion of the sequencing run. The FASTQ files were processed using SEISMIC-RNA (version 0.14.1) (Allan et al., 2024). Briefly, SEISMIC-RNA demultiplexes based on the barcode of a given reference, aligns reads to the known sequence, and quantifies the number of mutations on each read at each position. The function *wf* (for workflow) was executed including the option *-x* for 2 separate FASTQ files of paired-end reads; *--demult-on* for the second round of demultiplexing (a table with the barcode positions must be provided as well as a FASTA with the references, see Supplementary Table 1); *--sections-file* to annotate parts of the library sequence, including primers, flanking sequence, region of interest, and barcode; and *--export* for exporting a *json* file that can be provided to SEISMICgraph. Data were analyzed in SEISMICgraph. The filtered data for each plot were downloaded from SEISMICgraph and replotted in GraphPad Prism 10 to increase the resolution for publication.

## RESULTS

### SEISMICgraph: a web-based analysis tool

We developed SEISMICgraph (Figure 1 and Supplementary Figure 1) as a web-based application to address the need for rapid and simple visualization of chemical probing data, without requiring programming skills. SEISMICgraph consists of three main parts: (A) A front end user interface through which the user uploads data, makes selections, and requests plots; (B) A backend server that handles these requests and responds to them; (C) A python package (seismic-graph) that runs on the backend, processing uploads and generating plots.

SEISMICgraph uses data from chemical probing experiments to generate relevant plots. Users can upload data from multiple experiments at once, up to 32MB total. For datasets pre-processed with RNA Framework or SHAPEMapper 2, SEISMICgraph automatically converts into SEISMIC-RNA format on upload. For details on how the data is structured, transformed, and organized, refer to Supplementary Information section “Input data structure”.

#### User Interface

The user interface is separated into three main sections, (Figure 1B) Data Selections, Plot Selections, and the Plot Viewer.

In the Data Selections panel, users choose a subset of the uploaded data to plot. Certain plot types require only one row to be selected at a time, while others require two or more rows to be selected. Additional filtering options are available depending on the data selected, with more detailed information provided in the Supplementary Information section “Data Selection Options”.

Under Plot Selections, the user can select from several plots provided by SEISMICgraph. The Supplementary Information lists the plots that can be generated, and Figure 2 highlights examples of each plot type. The full details for additional plotting options are given in the Supplementary Information section “Plot Selection Options”.

Plot Viewer displays the rendered plot. The plot can be downloaded as an image or interactive HTML page. Additionally, the data can be downloaded as a CSV file for further analysis or to plot in ways not available in SEISMICgraph.

### SEISMICgraph Identifies Riboswitches and Pseudouridylation’s Impact on Function

We used SEISMICgraph to analyze DMS data for a designed library of RNA sequences to evaluate several predicted riboswitches, characterize several RNAs with well-established structures, and determine the effects of base modifications on the structure and function of these RNAs. Our library consisted of 81 RNAs, categorized as follows: 30 computationally-predicted purine riboswitches from various prokaryotic species (Mukherjee et al., 2019), 29 predicted Mg^2+^ riboswitches, and 22 other RNAs with well-characterized structures (Figure 3A). A complete list of sequences and their identities is provided in the Supplementary Table 1. Additionally, we provide detailed reports for all data and constructs in the Supplementary Dataset. We strongly encourage practitioners of chemical modification experiments to use and publish analogous summaries to make these studies more transparent and accessible to the scientific community.

#### Identification of Purine-Responsive Riboswitches

The designed library was transcribed *in vitro* and folded in the presence or absence of adenine. The samples were then subjected to DMS treatment followed by library-specific RT-PCR (Tomezsko et al., 2020) to detect conformational changes due to ligand binding (Figure 3A-C).

As a positive control, we first evaluated the adenosine deaminase (*add*) riboswitch from *Vibrio vulnificus*, which is known to alter its conformation upon adenine binding (Tian et al., 2018; Tomezsko et al., 2020). We demonstrated reproducibility in the DMS signal through two technical replicates, as shown in the **Mutation Fraction** plots of those replicates (Figure 3D and Supplementary Dataset “U1_Vibrio_vulnificus_0 and U2_Vibrio_vulnificus_0”) and the **Compare Mutation Profiles** scatterplot (Figure 3E, Pearson *R*^2^ = 0.98 and Supplementary Figure 2B, Pearson *R*^2^ = 0.99). In contrast, a comparison of the *add* riboswitch with and without adenine revealed differences, indicated by a much lower Pearson correlation coefficient relative to the replicates (Pearson *R*^2^ = 0.22; Figure 3F). These differences were also apparent from inspection of the mutation fraction across the sequence (Figure 3D compared to Supplementary Figure 2A) and the difference plot between these conditions (Figure 3G).

The ability to download data from SEISMICgraph for further analysis allows users to closely examine single nucleotides changes under different conditions. For the *add* riboswitch, we identified individual bases within the aptamer binding domain that substantially decreased in DMS reactivity when adenine was present (Figure 3H, I). The bases that were protected from DMS methylation upon ligand binding align with previous studies (Tian et al., 2018) and the hydrogen-bonding interactions enforced by adenine binding in the X-ray crystal structure of the ligand-bound state (Serganov et al., 2004). Such detailed analyses can enhance our understanding of the structural transformations induced by ligand binding. Future experiments can expand this approach to study the effect of other metabolites as well as studying these riboswitches in cells.

Next, we analyzed our entire RNA library using SEISMICgraph to compare the DMS reactivity in the presence and absence of adenine (Figure 3A). To provide an overall metric for adenine-dependent changes, we compared Pearson *R*^2^ values between replicates and between experimental conditions (Figure 3B, C). Three putative purine riboswitches, in addition to the established *add* riboswitch control, exhibited markedly lower *R*^2^ values (<0.6) compared to all other sequences (Figure 3B, C). Thus, DMS-MaPseq chemical probing can be used to identify candidate riboswitches. In all three putative riboswitches, the adenine binding pocket is highly conserved (Supplementary Figure 4B-E), and a decrease in DMS reactivity is observed at these bases, akin to what is seen in the *add* riboswitch (Figure 3J-L). The putative riboswitch from *Shewanella piezotolerans* is found upstream of a gene encoding adenosine deaminase (Figure 3L), and the other two putative riboswitches (from *Thermotoga thermarum* and *Thermotoga lettingae*) are located upstream of operons involved in *de novo* IMP biosynthesis (Figure 3J, K). Thus, it is reasonable that the three identified riboswitches are authentic, although direct functional tests in cells are needed. Equally interesting are the RNAs with sequence homology (Supplementary Figure 3) that did not change conformation upon addition of adenine. These might adopt different, functional structures in cells or have evolved to bind different ligands or carry out different functions.

**Figure 4.**
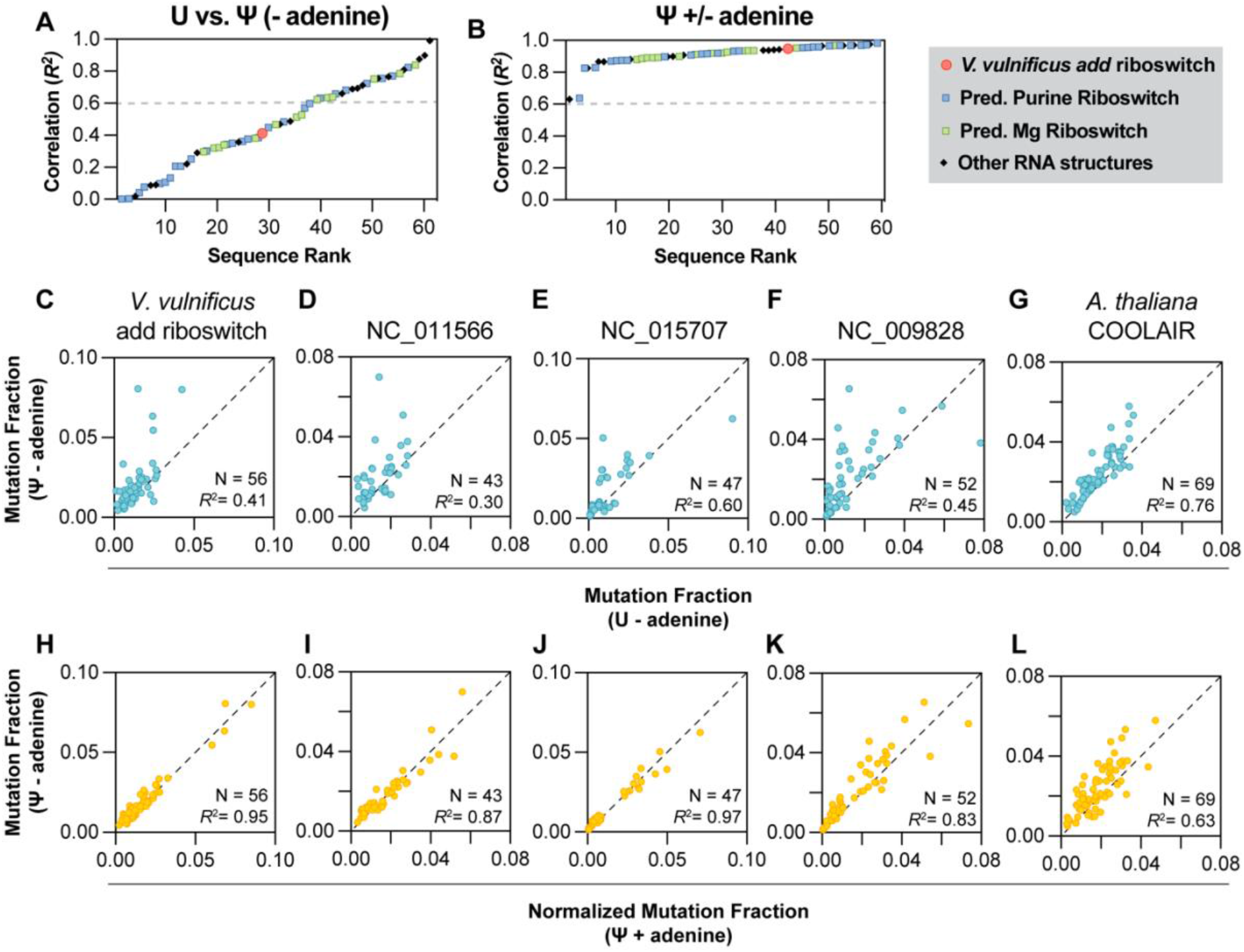
Effects of Pseudouridylation on RNA structure and riboswitching. (A) Ranked correlation of references compared to each other in samples containing all uridines or all pseudouridines. (B) Ranked correlation of references compared to each other in samples containing all pseudouridines in the presence and absence of 5 mM adenine. Each symbol represents one reference. (C-G) Comparison of the DMS signal of adenine responsive sequences from Figure 3 transcribed either with all uridines or all pseudouridines in the absence of adenine. Each turquoise circle represents one A or C base in the variable section. Neither sample was normalized. (H-L) Comparison of the DMS signal of adenine responsive sequences from Figure 4 transcribed with all pseudouridines in the presence or absence of adenine. Each yellow circle represents one A or C base in the variable section. The sample 2 (Ψ + Adenine) is normalized to sample 1 (Ψ – Adenine) as described in Supplementary Information. For (C-L), N represents the number of A and C bases. The Pearson *R*^2^ was calculated in SEISMICgraph.

Surprisingly, one RNA lacking homology to adenine riboswitches also appeared to adopt an altered conformation in the presence of adenine. A short sequence from the *Arabidopsis thaliana* gene COOLAIR also appeared to be responsive to adenine (Figure 3M and Supplementary Figure 5). While the 5’ domain of exon 1 from which this sequence was taken has several internal loops that might serve as a binding pocket for adenine (Yang et al., 2022), there is no observable pattern akin to that seen in the purine riboswitches. The shift in conformation may represent an accidental binding event (Peracchi et al., 1996) or an underlying yet-to-be-identified riboswitch function in a eukaryotic RNA. Regardless, the ability of RNAs to associate with nucleobases and other small ligands has implications for evolution in an RNA world and for potential modern-day RNA functions (Benner et al., 1989; Peracchi et al., 1996).

The “other” RNA structures included in the library primarily serve as controls, allowing us to confirm that these structures can fold similarly to previous reports and to ensure that the addition of adenine does not affect the chemical probing experiment in the absence of binding. For example, the frameshifting stimulation element (FSE) from HIV has a mutational profile consistent with stem-loop formation in the absence of adenine (Supplementary Figure 6A, B). This pattern remains unchanged in presence of adenine (Supplementary Figure 6C, D).

Putative Mg-sensing riboswitch structures have a motif or junction that could act as the binding pocket for Mg^2+^, similar to those found in known and putative purine riboswitches. We determined if sequences predicted to shift in response to Mg^2+^ also change conformations in the presence of adenine. Of the 15 sequences that passed quality control filters, none showed an observable conformational change (all Pearson *R*^2^ > 0.6, Figure 3C) at the working conditions (300 mM Na-cacodylate, 6 mM MgCl_2_). For example, sequence NC_0009792 appears to be folded in the absence of adenine (Supplementary Figure 7A, B), and its conformation does not change in the presence of adenine (Supplementary Figure 7C, D). Future experiments could measure the thermodynamics of folding by varying the Mg^2+^ concentration or temperature, using SEISMICgraph to analyze the results.

##### Pseudouridylation Prevents Conformational Switching Upon the Addition of Adenine

Pseudouridine (Ψ) and N1-methylpseudouridine (m1Ψ) are naturally occurring post-transcriptional modifications used widely in RNA therapeutics due to their ability to evade immune recognition and increase translation (Morais et al., 2021). Since pseudouridylation has previously been shown to exert both stabilizing and destabilizing effects on RNA structure (Davis, 1995; Meroueh et al., 2000; Vögele et al., 2023), we examined the functional consequences these modifications would have on the purine riboswitches analysed above. The procedure was repeated as before, but with uridine (U) replaced with either Ψ or m1Ψ during *in vitro* transcription of the library (Figure 4 and Supplementary Figures 8 and 9).

Incorporation of Ψ resulted in structural changes in more than half of the library sequences passing quality control filters (37 of 60; Figure 4A). These changes indicate broad structural transformations, with decreasing levels of change highlighted in Supplementary Figure 8. Similar DMS patterns, indicative of structural re-arrangements, were also observed in sequences modified with m1Ψ, detailed in Supplementary Figure 9B (17 of 59). These structural effects reflect the chemical properties of Ψ and m1Ψ, which both contain a C5-C1′ bond that augments base-pairing, base-stacking, and duplex stability, likely reducing the energy cost of non-native structures (Davis, 1995; Parr et al., 2020). While the DMS reactivities in Ψ and m1Ψ modifications generally demonstrated some correlation, indicating similarities in their structural impacts, the extent of these similarities varied depending on the sequence (Supplementary Figure 9C-D). This variability highlights that Ψ and m1Ψ exhibit distinct behaviours in different sequences, likely due to the fact that Ψ has an extra hydrogen bond donor at N1, which is absent in m1Ψ, and is therefore capable of wobble base-pairing with G, U, or C in a duplex (Kierzek et al., 2014). The differences are further suggested by the even more drastic structural changes seen upon incorporation of Ψ compared to m1Ψ. Notably, the introduction of either modification prevented conformational changes in all four adenine-sensing riboswitches and the COOLAIR structure (Figures 4 C-L). Given that Ψ has been shown to exert context-dependent destabilizing effects in functional RNAs, as in the *E. coli* 23S rRNA and the neomycin-sensing riboswitch (Meroueh et al., 2000; Vögele et al., 2023), modifications in the U-rich aptamer domain of the *add* riboswitch would likely also disrupt tertiary contacts crucial for native folding, thereby eliminating adenine-sensing and functional switching of the pseudouridylated riboswitches.

## DISCUSSION

We developed SEISMICgraph to streamline and standardize the analysis of chemical probing data. This tool provides various plotting and filtering options that enable rapid assessment and quantitative evaluation of experimental samples. It facilitates a broad overview by allowing researchers to examine multiple gene references simultaneously while also supporting detailed analysis by focusing on specific sequences. Additionally, SEISMICgraph allows for fast and thorough comparisons across different samples, experimental conditions, and sequences, making it a versatile and efficient resource for RNA structure research.

We confirmed three key findings using SEISMICgraph. First, we validated the structural changes in a well-documented adenine riboswitch. Second, we characterized the structures of a library of 81 sequences including 30 of putative purine riboswitches and found that three exhibit changes in response to adenine. Third, we demonstrated that introducing base modifications results in defects in structural switching. Notably, Ψ and m1Ψ modifications prevent the adenine-sensing function of the *add* riboswitch and the three other purine-responsive riboswitches.

These findings present a challenge for RNA therapeutics, as RNA modifications are often used to avoid host immune responses but may interfere with essential functions, as shown here. Chemical probing techniques such as DMS and SHAPE can help identify when and where these interferences occur. SEISMICgraph can be used to visualize and interpret these results, providing valuable insights for designing functional structures and developing RNA therapeutics.

## DATA AVAILABILITY

SEISMICgraph is available at https://seismicrna.org. Code is available at https://rouskinlab.github.io/seismic-graph/. All data collected in this study can be found at the Sequence Read Archive, accession PRJNA1165154.

## SUPPLEMENTARY DATA

Supplementary Data are available at NAR online.

## AUTHOR CONTRIBUTIONS

Federico Fuchs Wightman: Conceptualization, Library design, Validation, Formal analysis, Methodology, Writing-original draft. Yves J. Martin des Taillades: SEISMICgraph library development, Webapp UX/UI design, Writing-review & editing. Grant Yang: Conceptualization, Validation, Formal analysis, Methodology, Writing-review & editing. Casper L’Esperance-Kerckhoff: SEISMICgraph library implementation, Webapp UX/UI design and development. Scott Grote: Conceptualization and Library design. Matthew F. Allan: SEISMIC development and adaptation to SEISMICgraph. Daniel Herschlag: Conceptualization and Writing-review & editing. Silvi Rouskin: Conceptualization, Formal analysis, Writing-original draft. Lauren D. Hagler: Conceptualization, Library design, Methodology, Formal analysis, Writing-original draft.

## ACKNOWLEDGEMENTS

We thank the members of the Rouskin and Herschlag labs for the general discussion of the manuscript and their support.

