## Supplementary Information for "SEISMICgraph: a web-based tool for RNA structure data visualization"

### I. Input data structure

#### SEISMIC-RNA

The output of SEISMIC-RNA can be stored in a JSON format by using the option --export while pre-processing the sequencing data (Allan et al., 2024). By providing “samples'' and “sections” files using -S and -s respectively, the data output will be organized into a four-level hierarchical structure: sample, reference, section, and cluster. Note that while metadata at all levels is not necessary to run SEISMICgraph, defining these parameters will greatly enhance the capabilities and utility of the web app. Each level contains metadata and instances of the next level. Metadata names begin with a '#' sign. If level A contains level B, level A is referred to as the outer level, and level B is referred to as the inner level.

- Sample: A sample is a single fastq or json file. The sample level contains metadata about the experimental conditions of an entire fastq or json file (e.g., sample name, buffer conditions, temperature, DMS concentration, reaction time, etc.) and a list of references.
- Reference: A reference is the name of a sequence (either gene name or name of a unique sequence in a library provided in the FASTA file). The reference level contains metadata about one specific sequence (e.g., number of aligned reads, etc.) and a list of its sections (if provided).
- Section: A section is an identifier used to define a particular region of a sequence, such as a common motif present in all sequences, a specific region of interest, or the full sequence. The section level contains metadata about one specific region (e.g., name, sequence, and secondary structure prediction using RNAstructure, etc.) and a list of its clusters.
- Cluster: A cluster is a grouping of certain reads based on their observation of commonly occurring mutations on single molecules and are used to determine patterns of reactivity consistent with alternative RNA secondary structures (Tomezsko et al., 2020). Clusters are determined by an expectation-maximization algorithm in SEISMIC-RNA/DREEM as previously described (Allan et al., 2024). The "population average" cluster includes all reads

of a given section. The cluster level contains the mutation fraction, base coverage, and other data for each cluster.

Please refer to the official documentation for more details on the input format [docs].

**SHAPEmapper 2**

SHAPEmapper 2 outputs a tab-delimited text file that is primarily utilized for the analysis of SHAPE data (Busan and Weeks, 2018). This file contains comprehensive data for the entire experiment, distinguishing between treatments such as SHAPE-modified, untreated, and denatured samples. For each sample, the file includes information on the read depth, the identity of the mutation, and mutation fraction at each nucleotide position within the sequence. The data can be then converted from SHAPEmapper 2 to SEISMIC-RNA output by splitting it into three distinct samples. To conform to the hierarchical structure of SEISMIC-RNA, samples are designated as {sample_name}{sample_type}, where "sample_name" is derived from the directory name and "sample_type" indicates whether the sample is modified, untreated, or denatured. The file name serves as the reference. By default, the section parameter is set to "full", and the cluster parameter is set to "pop_avg". For more details, please visit the documentation.

**RNA Framework**

RNA Framework (Incarnato et al., 2018) generates a tab-delimited text file containing the reference name (on the first line), the sequence (on the second line), the mutation fraction per position (on the third line), and the number of reads at each position (on the fourth line).

The information is then processed equivalently to the SEISMIC-RNA output. For more details, please visit the documentation.

**Backend database structure**

After loading, the provided data is organized into a relational database format, characterized by a tabular structure with rows and columns. Each row within this database represents the contents of a single cluster, augmented by the metadata from all corresponding outer levels. Identification of each row is facilitated through a unique combination of identifiers pertaining to the sample, construct, section, and cluster.

**Filtering options and filtering reset**

When a filter is applied at a certain level, the inner levels' filtering options are updated such that the user can only select filtered-in inner levels. If there is an update on an outer filter that makes an inner level filter obsolete, the inner level filter will then be removed.

**Filtering arrays**

Filtering can also be conducted based on the positions within the sequence, offering the following methodologies:

- By Index: Users input 1-indexed boundaries to delineate a subset of the sequence for display.
- By Unique Sub-sequence: Users provide a specific sequence. This sequence must be present within the sequences of the rows intended for display. The data to be displayed will correspond to the indexes of this subsequence within the selected sequence.
- By Base Type: Users have the option to filter based on the selection of nucleotide bases, namely A, C, G, and T.

<u>Normalization</u>

SEISMICgraph incorporates a normalization feature that precedes the comparison of samples. This normalization process involves dividing the data from each sample by the slope of the linear regression calculated between the data of the first sample and the data from each of the other samples. In other words, the slope between all the reactive bases in two samples is used to transform one sample to the other, where y_normalized = y * (1/slope). Normalization in this manner should increase the similarity of the best-fit line and the line of identity, but it should not change the Pearson correlation. Nevertheless, normalization should be used with caution since it can lead to the loss of subtle structural features, especially in regions with low reactivity.

Furthermore, different normalization methods can introduce biases, particularly if not carefully tailored to the specific dataset, potentially distorting conclusions.

Download data subset

A button on the interface allows the user to download either the plot image and/or the raw data in table format of the plot that is displayed on the screen.

### II. Data Selection Options

To display particular rows of data, users have the option to apply filters via the interface. Certain plot types require only one row to be selected at a time while others require two or more rows to be selected. The filtering criteria available are the following:

- Minimal Base Coverage: Rows are selected only if the minimal coverage value surpasses a specified threshold across all samples. The default threshold is set at 1000.
- Experimental Variable: Users can choose a metadata attribute associated with the sample and specify the values to include in the filter. For example, one might only want to plot samples performed at 37 °C.
- Library Attributes: Users can choose a metadata attribute associated with the reference and specify the values to include in the filter. For example, references can be grouped in the library by common features (i.e., point mutations within a given structure).
- Sample: A sample is defined as a single fastq or json file. Samples can be selected individually to look at a single reference or simultaneously to compare replicates, in vitro vs. in vivo conditions, or across a variable such as the concentration of a metabolite.
- Reference: A reference is the name of a sequence (either gene name or name of a unique sequence in a library). Generally, one will compare a given reference across conditions, though reference-to-reference comparisons can also be made.
- Section: A section is an identifier used to define a particular region of a sequence, such as a common motif present in all sequences, a specific region of interest, or the full sequence.
- Cluster: Refer to the SEISMIC-RNA section.

### III. Plot Selection Options

SEISMICgraph offers a wide variety of plots, from which users can toggle through:

- # Aligned Reads / Reference as Freq. Dist.: A histogram showing the number of aligned reads per reference sequence in bins (Figure 2B) for a single sample.
- Base Coverage: A bar plot depicting the number of reads at each position for a given reference and section (Figure 2C). Only one reference and section from one sample can be plotted at a time.
- Mutation per Read per Reference: A histogram showing the frequency distribution of mutations per read in bins (Figure 2D). One reference and section from one sample can be plotted at a time.
- Mutation Fraction: A bar plot showing the mutation fraction per position (a ratiometric value of the number of mutations at a given position over the total number of reads at that position, Figure 2E). A single reference and section from a single sample can be plotted at a time.
- Compare Mutation Profile: A scatterplot comparing the mutation fraction at each position between two different samples for a given reference and section (Figure 2F). For this plot, at least two samples need to be loaded, while a single reference and section is selected. SEISMICgraph automatically displays the line of identity (x = y) and calculates the best-fit line and RMSE from linear regression. Additionally, it calculates the Pearson correlation. Of note, unnormalized samples may exhibit differences between the best-fit line and the line of identity that can often be attributed to differences in the actual DMS concentration used in the experiment. Normalization is offered by SEISMICgraph, please see above.
- Mutation Fraction Identity: A stacked bar plot, quantifying the number of mutations at a given position and identifying the mutation from either DMS-MaPseq or SHAPE-MaPseq after reverse transcription (Figure 2G). Note that because random mutations are inserted at a modified position, it is expected that there should be a variety of mutations. If the mutations at a given position are consistently from the same base, one should consider

other factors unrelated to the chemical probing experiment. Only one reference and section for a given sample can be plotted at a time.

- Experimental Variable Across Samples: A line-plot showing the mutation fraction over a changing experimental variable at each position of a reference sequence (Figure 2H). Either multiple or single nucleotide positions can be selected to be plotted at a time. While the Compare Mutation Profile plot can only show two samples at a time, where the x-axis is sample 1 and the y-axis is sample 2, the Experimental Variable Across samples plots the changing variable on the x-axis and the mutation.
- Mutation Fraction Delta: A bar plot of the difference in mutation fraction between two different samples at each position (Figure 2I), where the difference is calculated as the mutation fraction of sample 1 minus the mutation fraction of sample 2. As for the Experimental Variable Across Samples plot, caution is advised when comparing two unnormalized samples.
- Correlation by Refs. between samples: A scatterplot showing the correlation of each reference sequence with itself between two different samples (Figure 2J). Each reference is ranked in increasing order of correlation.
- One Pager: this option provides many plots for a single reference of a single sample (one row) in one page (pop-ups need to be allowed for it to load correctly).

### IV. Download Options and examples

Both the plot as an image and the data in a formatted table can be downloaded directly from the interface of SEISMICgraph.

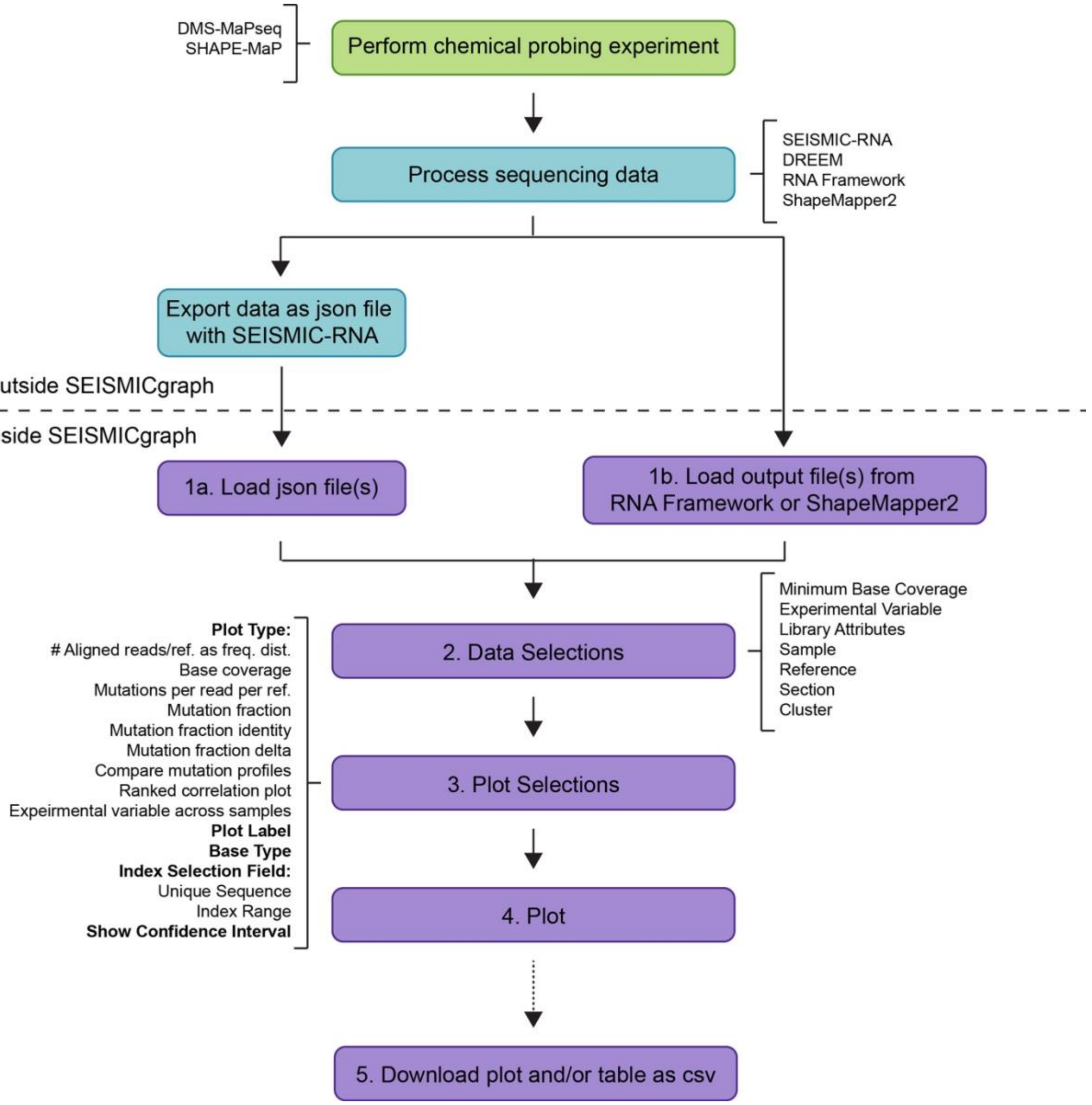


Supplementary Figure 1. Comprehensive workflow for SEISMICgraph. Steps prior to and using SEISMICgraph are delineated with options provided for every step.

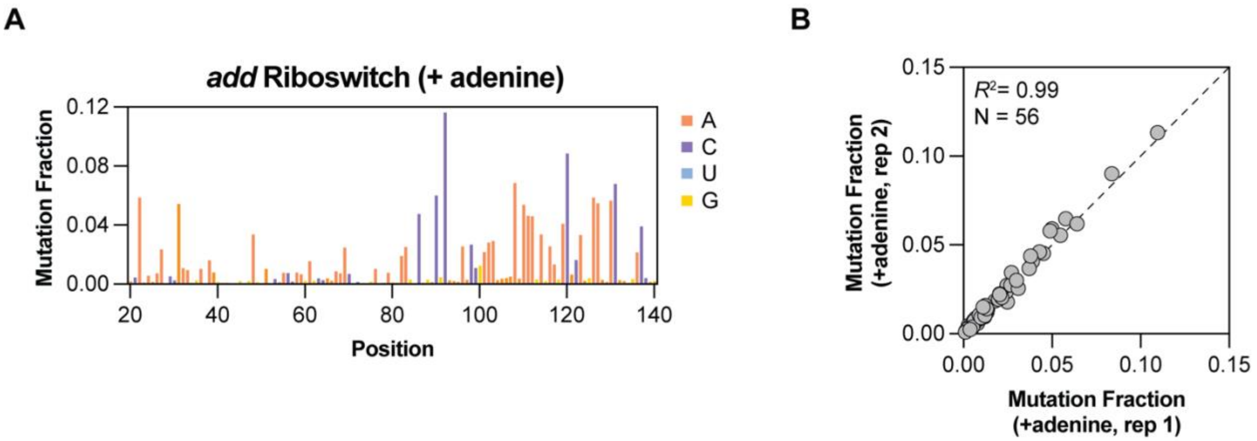


Supplementary Figure 2. Additional data for *add* riboswitch. (A) Mutation fraction per position for *add* riboswitch in the presence of 5 mM adenine. (B) Comparison of technical replicates with adenine.

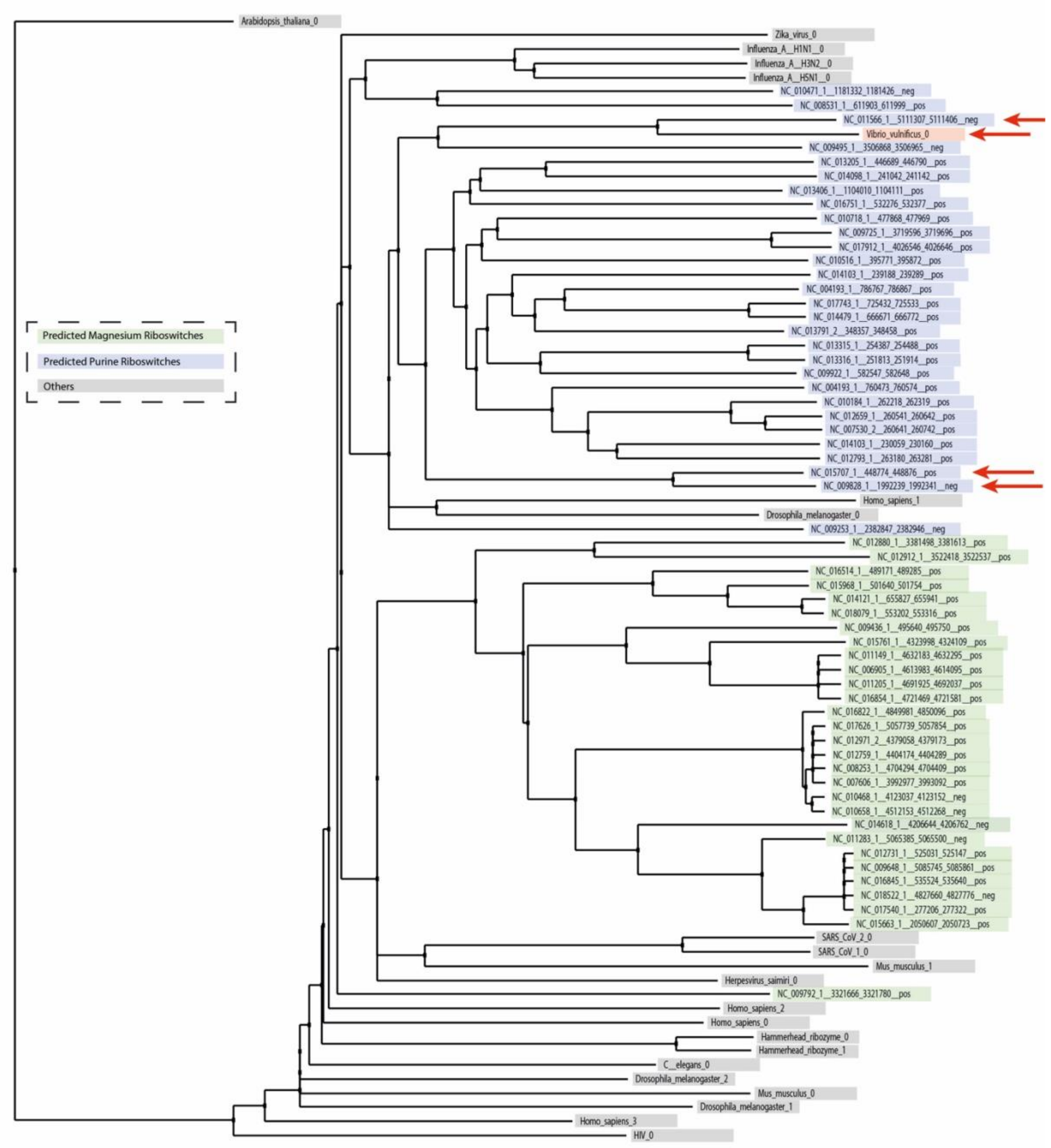


Supplementary Figure 3. Phylogenetic tree of RNA library. Predicted $Mg^{2+}$ riboswitches (green), predicted purine riboswitches (blue), and other structured RNAs (gray) are organized by sequence similarity as determined by Clustal Omega. Sequences that exhibited structural changes in response to adenine are marked with red arrows.

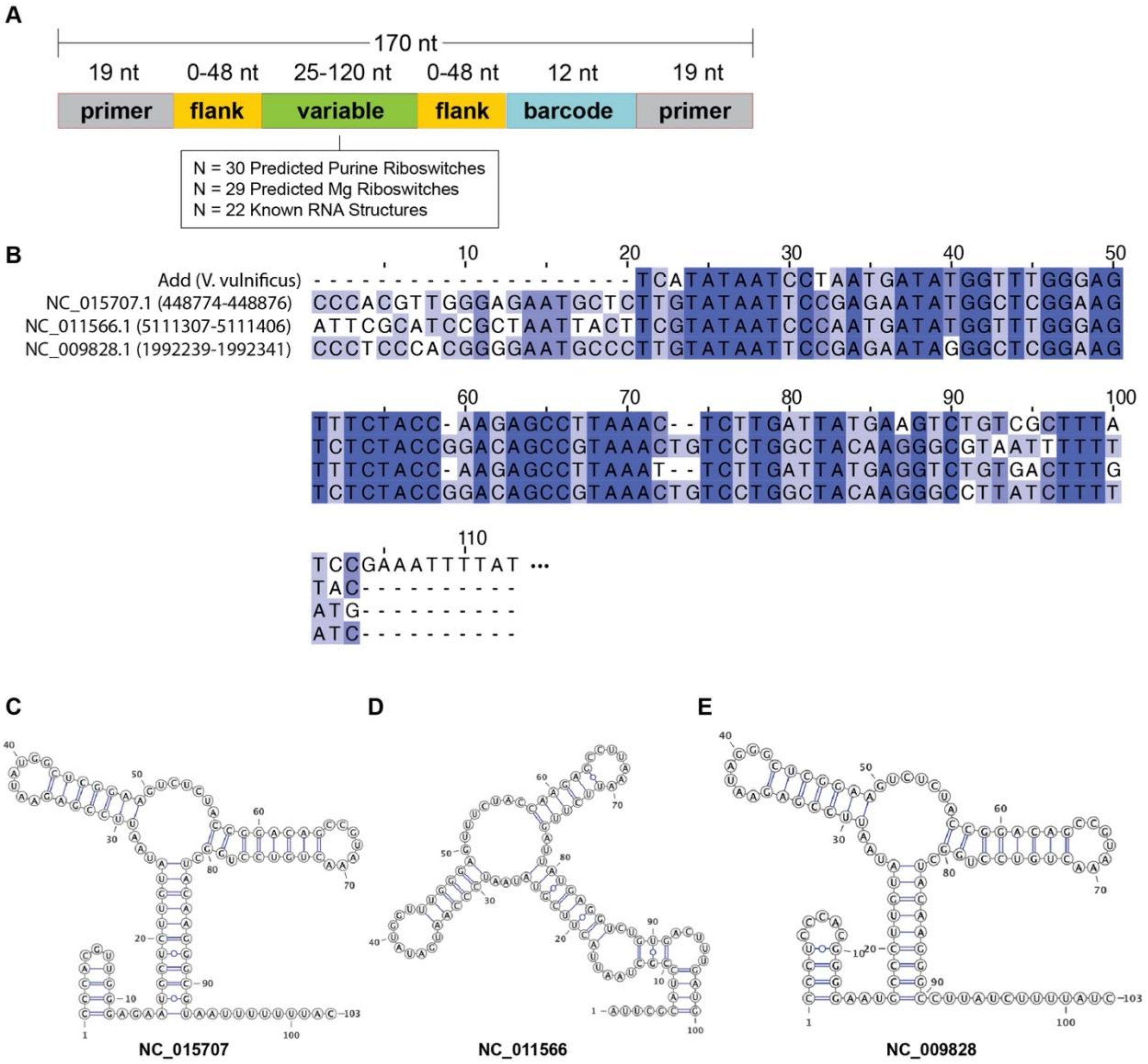


Supplementary Figure 4. Comparison of adenine-sensing riboswitches (A) Structural overview of the RNA library. (B) Sequence alignment of the *add* riboswitch and three putative riboswitches identified as responsive to adenine. (C-E) Predicted RNA structure of the three adenine-sensing riboswitches.

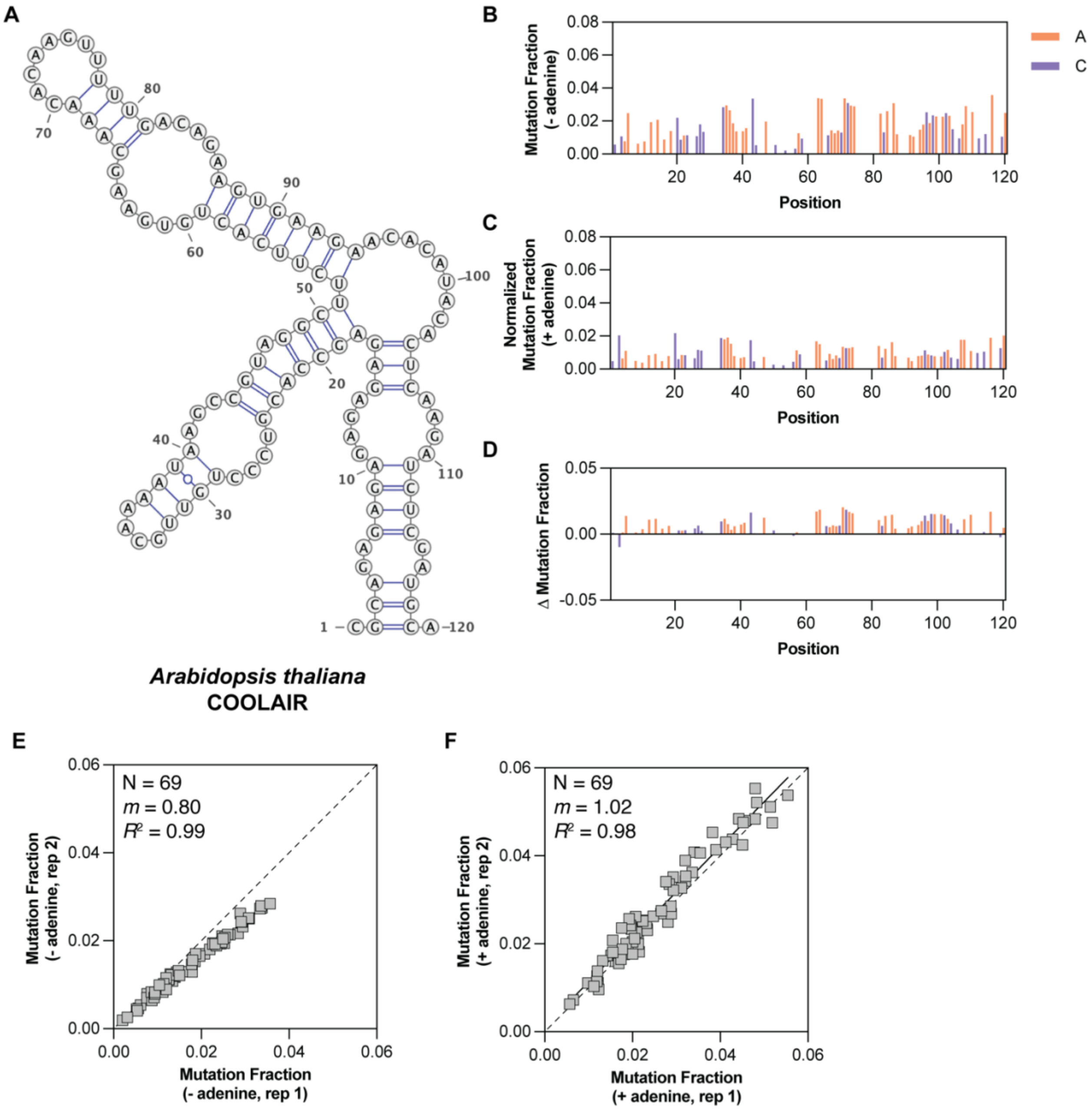


Supplementary Figure 5. Structural changes in the *Arabidopsis thaliana* COOLAIR transcript in response to adenine. (A) Structure model of library sequence from *A. thaliana* COOLAIR transcript. (B) Mutation fraction per position for COOLAIR in the absence of adenine. (C) Mutation fraction per position for COOLAIR in the presence of 5 mM adenine. (D) Difference in mutation fraction per position following addition of 5 mM adenine. (E-F) Comparison of replicate data in the absence and presence of 5 mM, respectively. Each grey square represents on reactive base. The black solid line represents te best-fit line from a linear regression, and the dashed line represents the line-of-identity. N is the number of reactive A and C bases, *m* is the slope of the best-fit line, and the $R^2$ is the Pearson correlation.

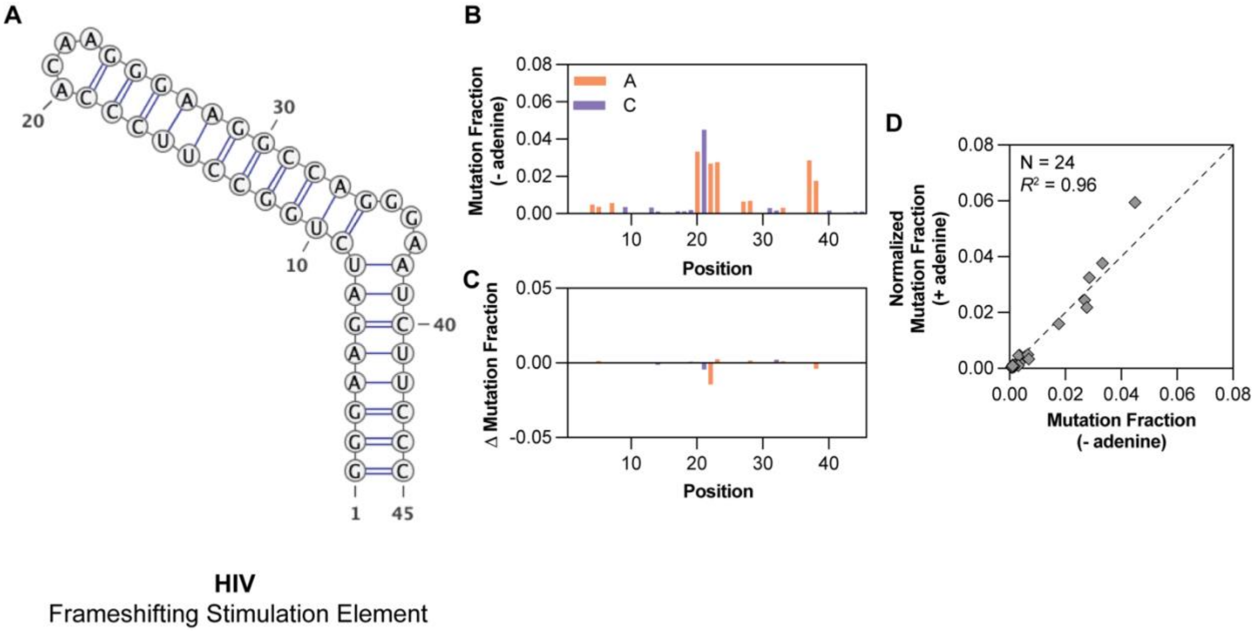


Supplementary Figure 6. Structural analysis of the HIV frameshifting stimulation element (FSE) in response to adenine. (A) Predicted RNA structure of the HIV FSE. (B) Mutation fraction per position in the absence of adenine. (C) Difference in mutation fraction per position following the addition of 5 mM adenine. (D) Correlation of mutation fractions with and without adenine.

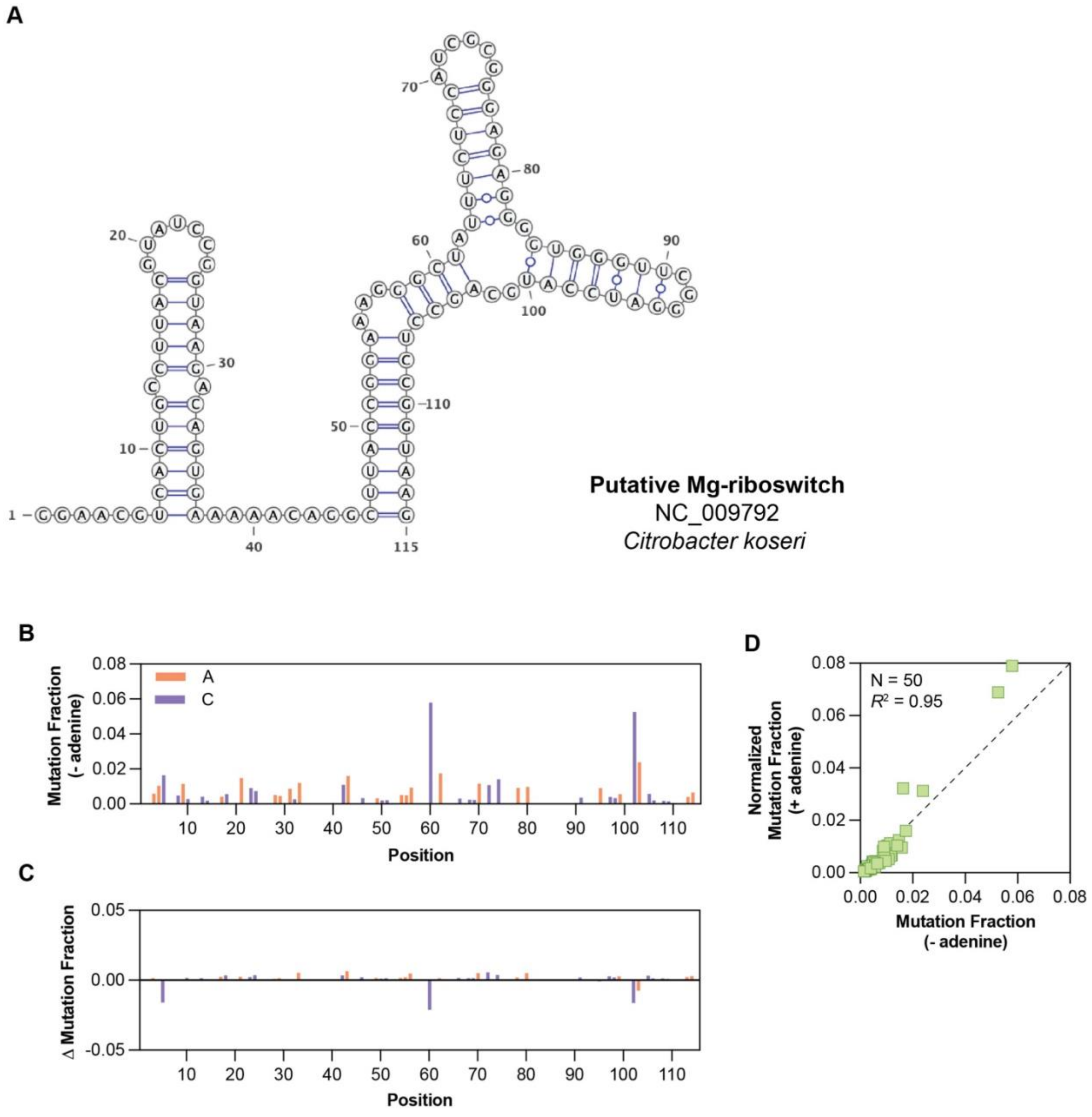


Supplementary Figure 7. Structural analysis of a putative $Mg^{2+}$ riboswitch in response to adenine. (A) Predicted RNA structure of predicted $Mg^{2+}$ riboswitch NC_009792. (B) Mutation fraction per position in the absence of adenine. (C) Difference in mutation fraction per position following the addition of 5 mM adenine. (D) Correlation of mutation fractions with and without adenine.

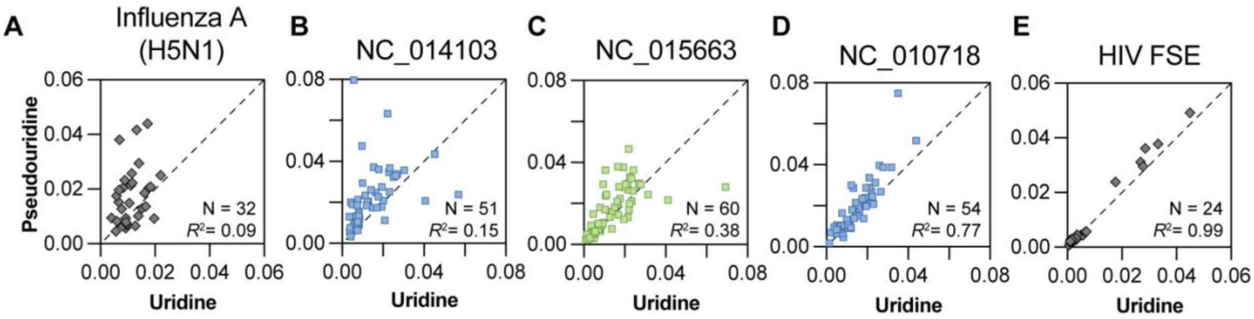


Supplementary Figure 8. Range of RNA structural perturbations caused by pseudouridylation. Increasing correlation between structures with all uridine or all pseudouridine is shown for (A) a sequence from Influenza A, (B) putative purine riboswitch NC_014103, (C) putative $Mg^{2+}$ riboswitch NC_015663, (D) putative purine riboswitch NC_010718, and (E) the HIV frameshifting stimulation element (FSE).

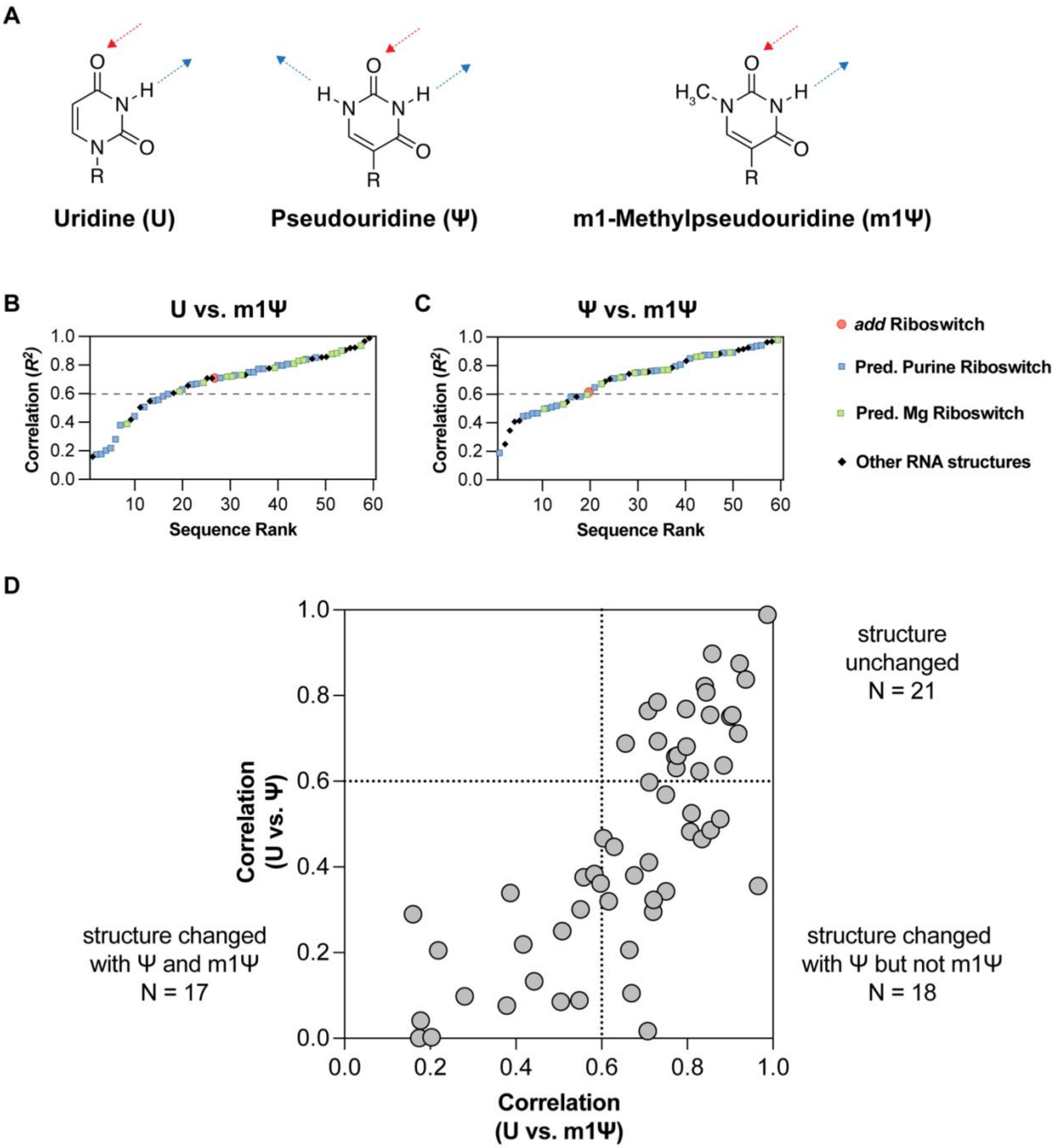


Supplementary Figure 9. Comparison of pseuduridylation and N1-methylpseudouridylation on RNA structure. (A) Chemical structures of uridine (U), pseudouridine (Ψ), and N1-methylpseudouridine (m1Ψ). (B) Ranked correlation of references in samples with all U or all m1Ψ. (C) Ranked correlation of references in samples with all Ψ or all m1Ψ. (D) Comparison of correlation changes for U vs. Ψ and U vs. m1Ψ. Number of structures that changed for each modification is labeled in each quadrant.
